# GeneScanner: profiling genetic variation across bacterial populations

**DOI:** 10.1101/2025.11.02.685864

**Authors:** Carolin M. Kobras, Seungwon Ko, Priyanshu S. Raikwar, Broncio Aguilar-Sanjuan, Keith A. Jolley, Samuel K. Sheppard

## Abstract

Rapid, low-cost genome sequencing has transformed microbiology, advancing efforts to link genetic and phenotypic variation. In laboratory settings, genome-wide functional screens of reference strains are revealing genes and mechanisms underlying important phenotypes. Simultaneously, population-scale comparative genomics describes the breadth of natural genetic and phenotypic diversity across lineages. These approaches provide complementary but contrasting information. Whereas laboratory studies offer causal understanding within simplified systems, population-level analyses capture ecological realism but are largely limited to detecting associations rather than cause-and-effect relationships. Linking laboratory-derived mutations to natural population variation remains challenging, particularly for researchers lacking bioinformatics expertise. Here, we present GeneScanner, a user-friendly tool that facilitates analysis of gene- and protein-level variation across large bacterial genome collections. GeneScanner detects genetic variation and amino acid substitutions in homologous sequence and supports genotype–phenotype association studies. By bridging laboratory functional genomics data with genomic diversity across natural populations, GeneScanner enhances functional interpretation of microbial variation in the real world, supporting research in antimicrobial resistance, virulence, and pathogen surveillance.

## Background

For decades, molecular microbiology has sought to understand the genetic basis of important bacterial traits including pathogenicity, antimicrobial resistance, and host interactions. Advances in sequencing technologies have transformed understanding of gene function, enabling the genome-wide analyses that define the field of functional genomics. For example, the principals of gene inactivation studies that compare the phenotypes of mutant and wild-type strains are now often augmented with genome-wide gene deletion [1–4] and transposon-insertion libraries [5–10], allowing rapid genotype-phenotypic screening under diverse selective conditions. Furthermore, in laboratory evolution experiments, where bacteria are subjected to selective pressures such as antibiotics or environmental stress [11–14], whole genome sequencing (WGS) is often used to compare progenitor and adapted isolates. While scientifically rigorous, these laboratory studies are often performed using a single laboratory adapted strain and may be limited in their ecological relevance. These approaches emphasize the importance of controlled experimental conditions and precise understanding of the ancestral and derived genetic background of laboratory strains, enabling targeted investigation of genotype-phenotype association. However, a laboratory settings may not fully reflect the complexity of natural environments where bacteria evolve [15].

The impact of WGS in a laboratory setting is mirrored in natural populations. Here the emphasis is often upon multi-strain comparative genomics for pathogen surveillance, phylogenetics and epidemiology. Millions of bacterial sequences are now freely available in public repositories [16,17] and curated databases [18–20], with metadata such as isolation source, place and time. This has transformed understanding of microbial genomics in the ‘wild’ with approaches including genome-wide association studies (GWAS) and covariation analyses revealing genetic variation underlying phenotypes linked to host adaptation [21–24], biofilm and virulence [25–30], and antibiotic resistance [31–34]. However, while some of the limitations of laboratory studies are overcome by analysing multiple strains in natural populations, highly relevant population genomics studies, lack the control and precision of laboratory functional genomics. This typically limits genotype-phenotype inference to association rather than cause-and-effect relationships.

Next-generation microbiology brings the rigour of the laboratory together with the relevance of population-wide studies and has major potential for understanding gene function in natural systems [35]. For example, studies of bacterial pathogen including *Campylobacter* [23] and *Streptococcus pneumoniae* [36] confirm laboratory observations in natural isolate populations allowing accurate analysis of functional genomics at a population scale. While database resources and isolate genome collections are freely available for comparable analyses of other pathogen species, analysis pipelines are often aimed at bioinformaticians rather than laboratory microbiologists. This can require the installation of specific operating systems and environments and command-line experience for single nucleotide polymorphism (SNP) calling [37–41].

To address this challenge, we developed GeneScanner, a user-friendly tool that enables researchers to explore genetic variation in specific genes or proteins across large datasets from natural bacterial populations. Implemented as both a stand-alone script on GitHub and a web-plugin available through PubMLST [20], GeneScanner simplifies the comparison of genetic features across strains and the identification of variants linked to phenotypic traits such as antibiotic resistance or pathogenicity. By providing an intuitive interface, it allows users without extensive bioinformatics expertise to analyse large datasets, contextualizing laboratory-observed genotypes within real-world bacterial diversity. In this study, we analyse synthetic data and three case studies across different bacterial species and phenotypes. We demonstrate how GeneScanner detects nucleotide and protein-level variation associated with specific traits, underscoring its broad applicability in microbial genomics, from antibiotic resistance studies to pathogen surveillance and laboratory evolution experiments.

## Results

### GeneScanner is a user-friendly tool for exploring genetic variation in bacterial populations

GeneScanner is a Python-based workflow, available on Github (https://github.com/Sheppard-Lab/GeneScanner), that accepts pre-aligned nucleotide or protein FASTA files. It screens every alignment column for sequence variation and collates the results into a multi-sheet workbook in .xlsx format. Even without bioinformatics background, GeneScanner can be used as a plugin on PubMLST (https://bigsdb.readthedocs.io/en/latest/data_analysis/genescanner.html) [20], supporting isolate selection and input file generation directly from organism-specific databases, and eliminating the need for local installation or command-line operation.

GeneScanner operates in four sequential stages (Fig. 1). In stage 1, the input alignment undergoes rigorous coverage and identity filtering, ensuring that only sequences meeting the predefined thresholds for ungapped alignment coverage and sequence identity to the reference are retained for subsequent analysis. In stage 2, the filtered sequences are analysed according to the user-selected mode (nucleotide or protein). In nucleotide mode, GeneScanner traverses the alignment both position-by-position and in codon triplets. Assuming no insertion or deletion events, each translated codon is compared with the corresponding reference codon to identify synonymous and non-synonymous substitutions, while insertions, deletions, and premature stop codons are recorded separately. In protein mode, the algorithm examines each aligned amino-acid column to quantify substitutions, gaps, and stop codons, and to calculate both mutation frequency and the remaining number of sequences following the stop codons. When the user requests protein analysis using nucleotide input, it automatically translates and realigns the sequences before proceeding as if native protein data had been provided.

**Figure 1.**
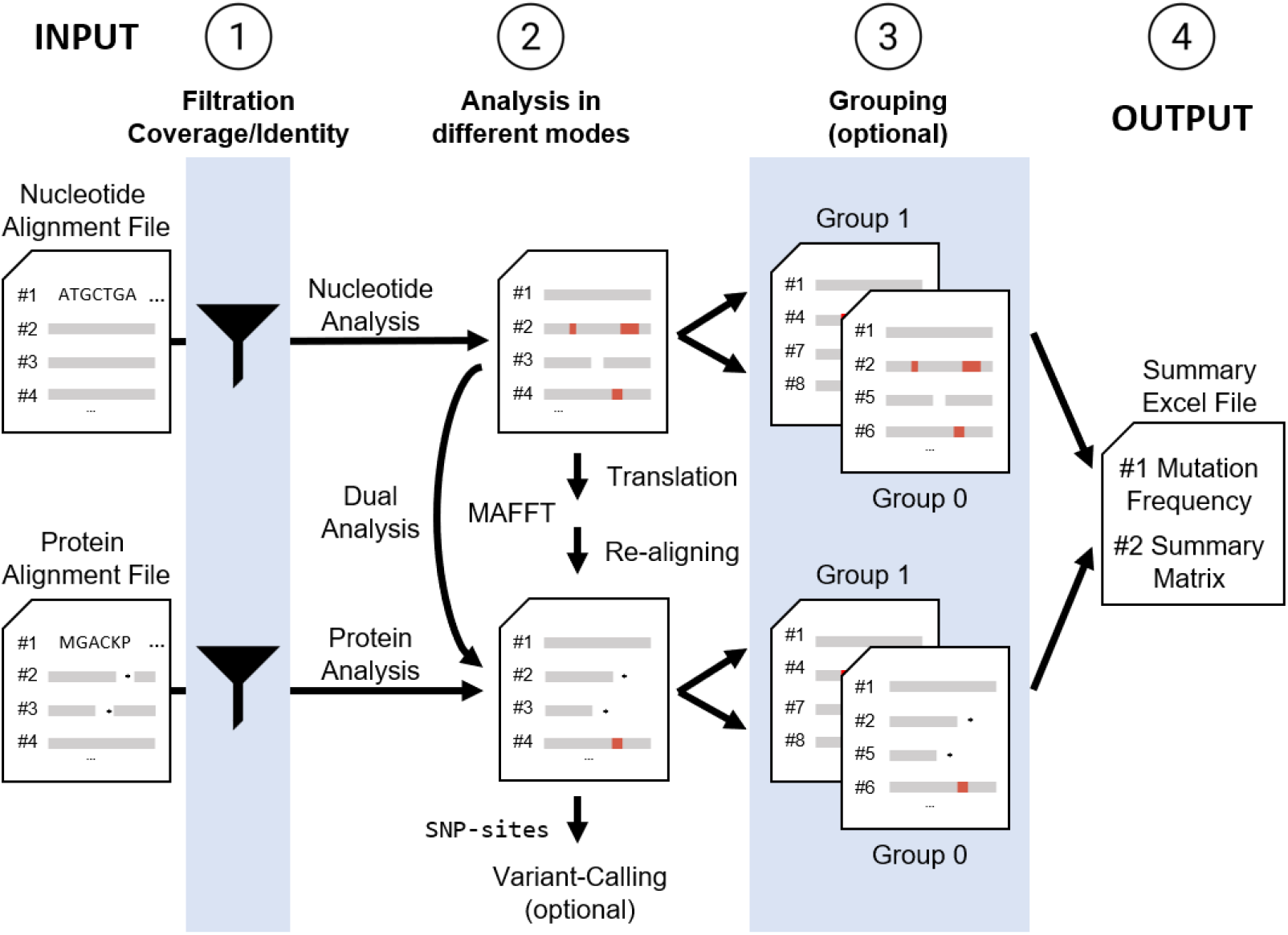
The GeneScanner workflow consists of four distinct steps. Schematic illustration of the GeneScanner workflow: (1) input parsing and quality filtering of alignment files (input); (2) codon-aware nucleotide or residue-aware protein variant calling; (3) optional per-group reanalysis; (4) output: xlsx format report generation with analysis, matrix, and summary worksheets. External tools MAFFT and snp-sites are invoked when translation, realignment or VCF output are requested.

In stage 3, GeneScanner can optionally perform grouping analyses when the user defines categories for sequences. This categorization helps subdivide the population, for example by phenotype or other defining traits. The program partitions the sequence dataset according to these categories and re-runs the full analytical pipeline for each subset. In stage 4, GeneScanner generates comprehensive nucleotide and amino acid mutation analysis spreadsheets that include detailed mutation categories. It also produces sparse mutation matrices in both nucleotide and protein modes, in which rows represent isolates, columns correspond to alignment positions, and each filled cell contains the variant symbol when it differs from the reference. Each matrix is written to a dedicated worksheet to facilitate downstream visualization and inspection.

GeneScanner also calculates and exports summary statistics to a dedicated worksheet. For nucleotide analyses, the summary includes the total alignment length, the number of mutated sites, positions with mutation frequencies exceeding 20%, and cumulative counts of synonymous and non-synonymous substitutions, and insertions, deletions, and stop-codons. For protein analyses, the corresponding statistics include substitutions, insertions, deletions, stop codons, and position-wise mutation frequencies. When multiple isolate groups are defined, GeneScanner generates separate worksheets and mutation matrices for each group, ensuring consistent column definitions and analytical parameters across all outputs. If distinct reference sequences are specified for individual groups, the program automatically reassigns the appropriate reference before performing variant calling.

### Benchmarking GeneScanner using a controlled synthetic dataset

To demonstrate the functionality of GeneScanner, we first tested it on a controlled synthetic dataset of 100 300bp sequences in which the locations, types, and frequencies of mutations were explicitly designed (Supplementary file 1). The dataset reproduced the intended patterns (Fig. 2A+B), and GeneScanner successfully captured variation at both nucleotide and protein levels (Supplementary File 2). At the nucleotide level, 20% and 30% synonymous variants occurred at positions 12 and 147, respectively, while 20% and 30% nonsynonymous variants appeared at positions 46 and 181, alongside a 10% nonsense mutation at position 210. When translated, synonymous sites produced no residue changes, whereas nonsynonymous mutations manifested as 20% Q16E and 30% S61P substitutions. The stop-gain mutation was correctly assigned to amino acid position 70, with GeneScanner distinguishing it from other nonsynonymous substitutions.

**Figure 2.**
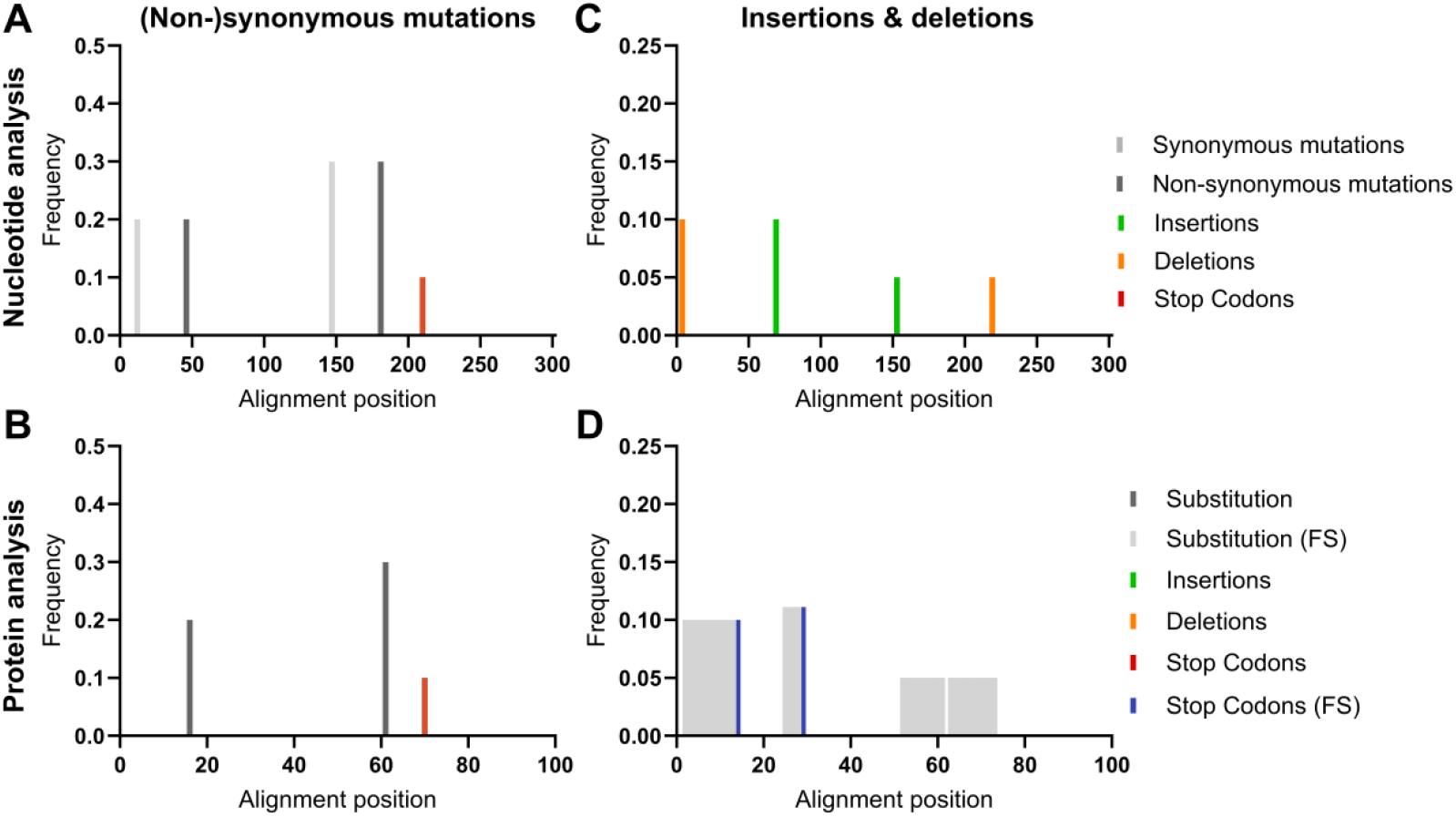
GeneScanner can reliably detect sequence variation on a nucleotide and protein level. Analysis of a synthetic dataset demonstrates and validates the functionality of GeneScanner. The tool successfully captures synonymous and non-synonymous mutations including those leading to stop codons at both nucleotide (A) and protein (B) levels. Insertions and deletions are recognised as events by nucleotide analysis (C), and as frameshifts (FS), premature stop codons or frame-restoration, where applicable, at the protein level (D).

The correct detection of insertion and deletion (indel) events further highlighted the flexibility of the workflow (Fig. 2C+D, Supplementary files 3+4). A single-nucleotide deletion at position 4 produced a frameshift with an early stop at amino acid position 14 (10%), and a single-nucleotide insertion at position 69 generated a frameshifted protein truncated at position 29 (10%). In the frame-restoration scenario, a paired insertion at position 153 (5%) and deletion at position 219 (5%) created a transient frameshift with amino acid substitutions between residues 52–73, after which the original reading frame was recovered. These results illustrate how GeneScanner’s dual nucleotide-protein mode can trace both the disruption and recovery of coding frames, preserving alignment quality while recording complex mutational events.

### GeneScanner validation of in silico sequence evolution across two populations

To further validate the functionality of GeneScanner on datasets with unknown outcomes, and to test its grouping feature for analysing distinct subpopulations, we simulated evolution with differential selection. We created two populations of 5,000 bacterial genomes, starting with a random 300 bp coding sequence. Mutations occurring in a specific region (positions 100–130) were set to have opposite fitness effects in the two populations, while mutations elsewhere were neutral. After 1,000 generations, we sampled 500 isolates from each population (p1 and p2) and ran GeneScanner (Fig. 3A, Supplementary files 5+6). The imposed selection produced clear divergence within the targeted 100-130 bp window (Fig. 3B+C). Using the original sequence before simulation as a reference, subpopulation p2 exhibited minor variation across the locus, consistent with neutral dynamics outside the selected window. In contrast, subpopulation p1 displayed a concentration of both synonymous and nonsynonymous differences restricted to positions 100-130, reflecting the impact of opposing selection pressures. Outside of the region under simulated, both p1 and p2 showed similar mutational patterns. GeneScanner successfully recapitulated these differences at nucleotide levels, confirming the tracking of mutational events across divergent evolutionary trajectories.

**Figure 3.**
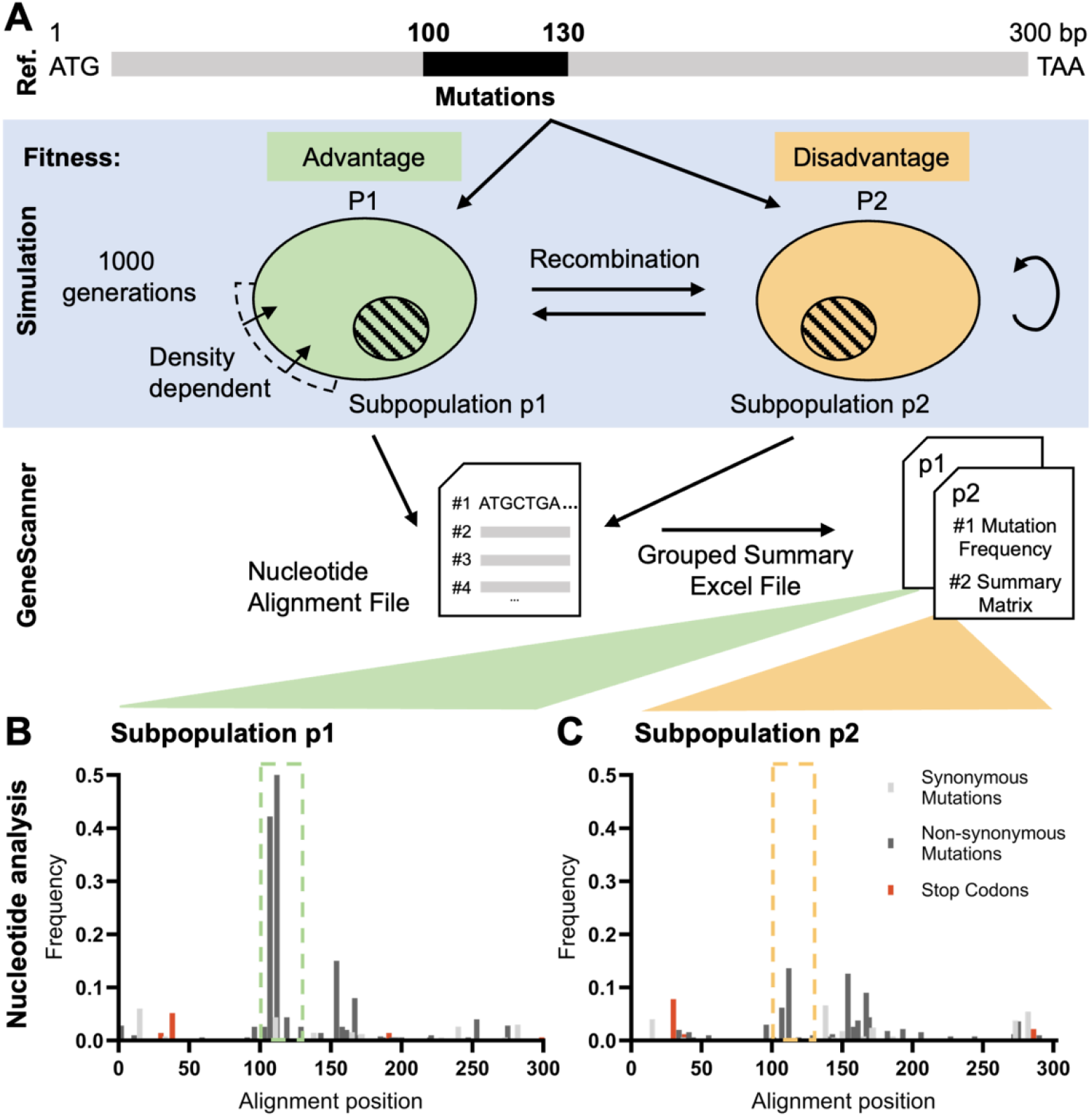
GeneScanner reliably detects sequence variation from *in vitro* sequence evolution under differential selection. A: Simulated sequence evolution produced divergent outcomes between two subpopulations exposed to opposing selection pressures. B+C: Using the original reference sequence, GeneScanner detected only scattered minor variation across the locus in p2, whereas p1 exhibited concentrated synonymous and nonsynonymous substitutions within the 100-130 bp window. These differences captured for both subpopulations, demonstrating the tool’s ability to resolve selection-driven divergence.

Together, these analyses demonstrate the utility of GeneScanner for detecting, classifying, and contextualizing intra-genic variation in bacterial populations. Using a controlled synthetic dataset, the workflow reproduced expected mutation patterns, and in simulations of *in silico* sequence evolution across different subpopulations, it accurately traced selection-driven divergence. This proof-of-concept evaluation established the four-stage GeneScanner pipeline - from alignment parsing to mutation classification and summary statistic generation. Building on this foundation, GeneScanner was used in three case studies spanning distinct bacterial species, phenotypes, and research contexts.

### Case study 1: Genetic variation of DNA replication genes is closely linked to ciprofloxacin resistance

Antibiotic resistance is a major global health concern, with *Escherichia coli* being among the most problematic pathogens [42]. The fluoroquinolone antibiotic ciprofloxacin is a commonly used treatment that targets essential enzymes involved in bacterial DNA replication, specifically DNA gyrase and topoisomerase IV [43,44]. However, *E. coli* can rapidly develop resistance to ciprofloxacin through mutations in genes linked to DNA replication [45]. In this case study, we used GeneScanner to highlight the most common amino acid substitutions associated with ciprofloxacin resistance in a natural population of *E. coli* isolates (n = 1509). Using GeneScanner, we analysed the DNA gyrase subunits A and B (GyrA, GyrB), and the topoisomerase IV subunits A and B (ParC and ParE) [45], across ciprofloxacin susceptible (n=1229) and resistant (n=280) isolates. To ensure accurate detection of group-specific variations, GeneScanner performs nucleotide and amino acid alignments before the populations are divided into susceptible and resistant groups, allowing the tool to accommodate insertions that may be unique to one group, and the resulting difference in alignment. Comparing the ProteinAnalysis sheets between groups, we identified several well-known amino acid changes at high frequencies in the resistant population. For example, over 97% of ciprofloxacin resistant isolates had GyrA substitutions at S83, and approximately 92% at D87 [45]. However, approximately 8% of the susceptible isolates also carried the S83 substitution, indicating that this change on its own may only confer small increases in resistance. Similarly, known resistance-associated ParC substitutions, such as S57, S80 and E84, were clearly overrepresented in our resistant population, suggesting that most isolates carried several resistance mutations. We further identified ParE^I529L^ in 153 resistant isolates, a substitution associated previously associated with *E. coli* ST131 [46]. Interestingly, we also identified ParC substitution A192V in 151 of these isolates by comparing the ProteinMatrix, indicating co-variation between these otherwise well-conserved genes, and further highlighting the utility of GeneScanner.

### Case study 2: Vitamin B5 biosynthesis genes are conserved in Campylobacter jejuni isolated from cattle

*Campylobacter jejuni* is a leading cause of bacterial gastroenteritis in humans, most often transmitted through the consumption of contaminated food, especially poultry [47–49]. *C. jejuni* colonises multiple hosts, including chickens, cattle, and wild birds, and distinct lineages frequently display host preference, consistent with niche specialisation through adaptation. One such adaptation involves the pantothenate (vitamin B5) biosynthesis pathway encoded by the *panBCD* locus. The first formal bacterial genome-wide association study (GWAS) [21] revealed that host association signals arose from two major forms of genetic variation. First, genes within the primary host-associated region were more commonly absent from chicken isolates. Second, when present, their sequences differed between cattle and chicken isolates, with cattle alleles exhibiting reduced homologous sequence variation – consistent with gene conservation. As a result, GWAS revealed host-association genes and alleles.

We used GeneScanner in a second case study to validate these findings in a larger collection of *C. jejuni* ST-45 complex isolates, comprising 277 from cattle and an equal number from chickens. Consistent with previous results, the *panBCD* locus was more common in cattle isolates (69%) than in those from chickens (60%), as indicated on the output sheet. Additionally, chicken-associated isolates generally exhibited a greater diversity of unique alleles for the *panBCD* locus. This pattern was confirmed by GeneScanner, which confirmed the elevated overall genetic variation among chicken isolates. For instance, although the number of unique *panB* alleles was similar between cattle and chicken isolates, the 826 nucleotide *panB* alignment contained 18 variable sites in isolates from cattle, whereas chicken isolates had 80. A similar trend was observed for *panC*, with 21 vs. 88 variable sites, with only *panD* showing more comparable values with 26 vs 35 variable sites in cattle and chicken isolates respectively. These statistics are readily accessible in the Summary Statistics tab of the output files.

Overall, GeneScanner analyses refined previous findings by pinpointing host-associated sites of genomic differentiation and conserved coding regions. This higher-resolution view confirmed the enrichment of the *panBCD* in cattle-associated lineages, while revealing markedly greater allelic diversity in chicken-associated lineages. Thus, GeneScanner strengthened the evidence for host-driven genomic signatures shaped by the interplay of metabolic capability, gene flow, and selective pressures in *C. jejuni* populations.

### Case study 3: Intergenic variation suggests strain-level modulation of biofilm expression

The third GeneScanner case study focused on cross-species comparison of a non-coding but significant regulatory region in staphylococci. The <u>i</u>nter<u>c</u>ellular <u>a</u>dhesion (*ica)* locus has an important role in biofilm formation in *Staphylococcus aureus* [50,51], contributing to persistent infections and treatment failure. The locus comprises *icaADBC* genes, which encode enzymes involved in polysaccharide intercellular adhesin (PIA) production, and the divergently transcribed repressor *icaR. icaADBC* expression is regulated by multiple elements beyond canonical transcription and translation initiation sites. Hence, the *icaR*_*icaA* intergenic region contains binding sites for several regulatory proteins, including the repressors IcaR [52], TcaR [53,54], and Rob [36], and the activator SarA [56]. The presence of the 163 nucleotide-long intergenic region is highly conserved across a diverse *S. aureus* isolate population [57], but despite nucleotide-resolution analysis of regulator binding sites in model strains, the genetic variation of the *icaR_icaA* intergenic region remains understudied.

GeneScanner nucleotide analysis revealed multiple loci with high levels of genetic variability within the *ica* promoter region, particularly near the start codons of *icaR* and *icaA*, as well as within the IcaR binding site (Fig. 6). Notably, we also observed variation within the Rob binding site, including modifications and complete loss of the TATTT motif (Fig. 6, orange box), which is essential for Rob recognition and binding. Such changes are likely to impair Rob-mediated repression, potentially leading to increased *ica* expression and enhanced biofilm formation. These patterns of variation suggest the potential for strain-specific modulation of *ica* operon activity and biofilm production. In contrast, several SarA binding sites appeared to be highly conserved across all *S. aureus* isolates analysed, indicating that SarA may serve as a consistent activator of *ica* expression across diverse genetic backgrounds.

Consistent with previous reports, the *ica* operon was absent in more distantly related staphylococcal species [50]. In contrast, 482 out of 1000 *S. epidermidis* isolates harboured the *ica* operon. However, GeneScanner summary statistics revealed only five variable sites within the *icaR_icaA* intergenic region, compared to 58 in *S. aureus*. This indicates a markedly higher degree of regulatory intergenic variation in *S. aureus*. Together, these findings demonstrate the utility of GeneScanner for interspecies comparison of non-coding regions, providing insights into the evolution of regulatory elements and their potential impact on phenotypic diversity.

## Conclusion

Extensive bacterial genome datasets provide major opportunities to study microbial diversity, evolution, and adaptation [58]. However, connecting mutations discovered in controlled laboratory experiments with the naturally occurring variation present in bacterial populations remains challenging [15,35]. This methodological disconnect is particularly pronounced for researchers without extensive bioinformatics expertise, as most available tools require complex computational skills and resources. GeneScanner is designed to bridge this gap by offering an accessible and powerful platform for analysing gene- and protein-level variation across large bacterial genome collections. The tool efficiently identifies both nucleotide variation and protein-altering amino acid changes in genes and proteins associated with phenotypic traits, including antibiotic resistance studies and pathogen surveillance. Demonstrating the functionality and broad applicability of GeneScanner, we applied it to designed and simulated datasets as well as three distinct bacterial datasets encompassing different species, phenotypes, and gene loci. In each case, the tool successfully identified nucleotide-level and protein-altering changes associated with relevant traits, illustrating GeneScanner’s utility as a comprehensive and versatile platform for investigating genetic variation across a wide range of bacterial systems, traits, and experimental contexts.

As with most genome analysis tools, the reliability of results depends heavily on data quality and sampling design. While GeneScanner includes quality control, poor assemblies, incomplete genomes, or incorrect annotations may introduce artefacts. Furthermore, biased sampling can distort allele frequencies, for example, overrepresentation of certain lineages can generate genotype-phenotype associations driven by population structure rather than true causality. To address this, analyses can incorporate population subsampling [28,59], linear mixed models [60,61], and phylogenetic approaches [62] to account for the clonal structure of the population. Additionally, using high-quality, well-annotated genomes and including metadata on isolation source, geographic origin, and collection date can help control for potential confounding factors.

Uncovering the structural and functional impacts of naturally occurring genetic variation will be essential for advancing our understanding of microbial adaptation and phenotype variation. GeneScanner provides a powerful starting point by efficiently identifying candidate SNPs and amino acid changes across large bacterial genome collections. Of course, understanding how these alterations influence protein conformation, stability, or activity will require additional analysis. Integrating structural bioinformatics methods - such as domain conservation analysis [63,64], structure–function prediction [65–67], and molecular dynamics [68,69] - alongside experimental validation could achieve this. Advancing microbiology in this direction will move future research beyond merely detecting variation, toward mechanistically explaining how it drives microbial phenotypes, maximizing the impact of bacterial genomic data on surveillance, outbreak response, and therapeutic development.

## Methods

### Technical specifications

All analyses were executed on PubMLST using the GeneScanner wrapper plugin [20]. Core software versions were Python 3.6+, Biopython 1.84 [70], pandas 2.2.2 [71], XlsxWriter 3.2.0 [72], MAFFT 7.490 [73], and snp-sites 2.5.1 [37]. Detailed requirement for each release version and all code are available in the public GitHub repository (https://github.com/Sheppard-Lab/GeneScanner).

### GeneScanner pipeline and analytical framework

Multiple-sequence alignment FASTA files are the primary input for GeneScanner. Two quality filters are applied relative to a designated reference sequence: a default minimum of 80% ungapped coverage across the alignment length and at least 80% pairwise identity. If requested, a VCF file is generated via snp-sites [37]. In nucleotide analysis mode, GeneScanner iterates through the alignment position-by-position while preserving codon phase relative to the reference, which can be user-defined or automatically set as the first sequence in the alignment. For each non-gap codon triplet, the translated codon is compared with the reference translation to classify synonymous and non-synonymous mutations. Insertions, deletions, and premature stop codons are recorded independently. When protein analysis mode is activated, GeneScanner first removes all the gaps within the original alignment file, translates in the user-specified frame and realigns peptides with MAFFT [73]. To prevent alignment disruption caused by early stop codons, the program temporarily fills post-stop-codon regions with reference residues, which are removed after realignment to maintain alignment quality.

Optional grouping analyses are triggered when the user provides a two-column CSV file specifying isolate names and their categories. GeneScanner partitions the dataset by category and re-runs the full analytical pipeline for each subset, producing parallel mutation analysis reports and mutation matrix worksheets that share column definitions but are restricted to their respective groups. When distinct reference sequences are supplied for individual groups, GeneScanner automatically reassigns the appropriate reference prior to variant calling. After all analyses are complete, GeneScanner compiles the results into a spreadsheet (.xlsx) by using pandas [71] and XlsxWriter [72]. All parameters are user-configurable via command-line flags and run metadata as well as quality-control filtering logs can be written to separate files.

### Synthetic sequence design for GeneScanner benchmarking

To create a dataset with explicitly designed mutations, we generated a 300 bp reference coding sequence and designed point mutations and short indels using custom Python code from the GeneScanner repository. To evaluate coding-effect categorization, we placed synonymous substitutions at alignment positions 12 and 147 with allele frequencies of 0.20 and 0.30, respectively; nonsynonymous substitutions at positions 46 and 181 with frequencies of 0.20 and 0.30, and a nonsense stop codon substitution at position 210 with frequency 0.10 to confirm that GeneScanner treats stop-gains distinctly from other variants. To assess frameshift handling, we designed two indel scenarios with single-nucleotide changes: (i) frameshift leading to premature termination via a deletion at nucleotide position 4 (10%) and an insertion at position 69 (10%) and (ii) a frame-restoring pair consisting of an insertion at position 153 (5%) followed by a downstream deletion at position 219 (5%) such that the reading frame is restored after a segment of shifted translation. All coordinates are based on the aligned nucleotide sequence and amino-acid coordinates refer to the translation of the reference.

### In silico sequence evolution simulation for GeneScanner validation

Sequence evolution of two subpopulations was simulated under opposing selection regimes using SLiM v5.0 [74] with nucleotide-based models and phylogeny data. The model script was generated applying the previous published code [75]. The simulated genome comprised a 300 bp random ancestral coding sequence. Mutations followed a symmetric Jukes-Cantor model with a per-site mutation rate of μ = 1 × 10^−6^ per generation. Recombination occurred at a rate of ρ = 1/3 × 10^−4^ per site per generation along the locus with 30bp as average recombination block length. At generation 1, we split the population into two panmictic subpopulations of 5000 haploid individuals each. Selection was restricted to a 31 bp window spanning nucleotide positions 100 - 130. Fitness was calculated by multiplying over sites and across the two populations. We assigned opposite selection coefficients in the two subpopulations: In subpopulation p1, mutations within the window were beneficial (s = + 0.005), whereas in subpopulation p2, the same mutations were deleterious (s = −0.005). Mutations outside the window mimicked neutral mutation with slightly deleterious fitness cost (s = −0.0001). Total fitness is calculated by 1 +∑ *s* compared to 1 at starting reference gene. To limit the extent of linkage hitchhiking, we chose an elevated recombination rate, as theory predicts that the influence of a selective sweep on linked diversity decreases with recombination distance relative to selection strength [76]. Density dependent fitness adjustment was added to keep the population size near the original 5,000 haploid genomes per population while it changes the sequence frequency within populations. Simulations were run for 1,000 generations under SLiM’s default non-Wright–Fisher framework. At the final generation, 500 sequences were sampled from each subpopulation. Sampled sequences were aligned using MAFFT, and mutational patterns were subsequently characterized using GeneScanner in dual nucleotide– protein mode.

### Case study 1: genetic variation underlying ciprofloxacin resistance in E. coli

The *E. coli* genome assemblies (n=1509, Supplementary file 7) from a previous study [77] were accessed on the PubMLST database [20]. Information on ciprofloxacin susceptibility of these isolates was taken from supplementary material. Nucleotide sequences of query genes (*gyrA, gyrB, parC* and *parE*) were downloaded from GenBank and used to query against isolate assemblies using blastn (BLAST 2.12.0+ with settings: word size: 20; reward: 2; penalty: −3; gapopen: 5; gapextend: 2). The identified matching sequences were extracted and aligned using MAFFT v7.505 with default parameters to create the nucleotide alignment input file. GeneScanner was run in ‘nucleotide + protein’ mode, selecting strain *E. coli* K12 MG1655 (PubMLST *Escherichia* ID: 1167) as reference for both groups. A CSV file for group 2 (resistant isolates) was created. Selected data were, as mutation frequency and normalised by the isolate number, visualised using GraphPad Prism (version 10.5.0)

### *Case study 2:* Conservation and diversity of vitamin B5 biosynthesis genes in *Campylobacter*

The genome assemblies of *Campylobacter jejuni* ST-45 complex isolates were accessed on the PubMLST database [20], using all available isolates resulting from the search term “cattle” (n=277), and randomly selecting an equal number of isolates containing the term “chicken” in the search field (Supplementary file 7). Nucleotide sequences of the *panBCD* locus intergenic were downloaded from GenBank and used to query against isolate assemblies using blastn (BLAST 2.12.0+ with settings: word size: 20; reward: 2; penalty: −3; gapopen: 5; gapextend: 2). The identified matching sequences were extracted and aligned using MAFFT v7.505 with default parameters to create the nucleotide alignment input file. GeneScanner was run in ‘nucleotide + protein’ mode, selecting strain *Campylobacter jejuni* NCTC11168 (PubMLST *Campylobacter jejuni/coli* ID: 48) as reference. A CSV file for group 2 (isolates from cattle) was created. The number of unique alleles was determined using the BIGSdb plugin GenomeComparator [78] using the same isolates as input and default settings. Selected data were, as mutation frequency and normalised by the isolate number, visualised using GraphPad Prism (version 10.5.0). Icons in Figure 5 were designed by PLANBstudio and sourced from flaticon.com.

**Figure 4.**
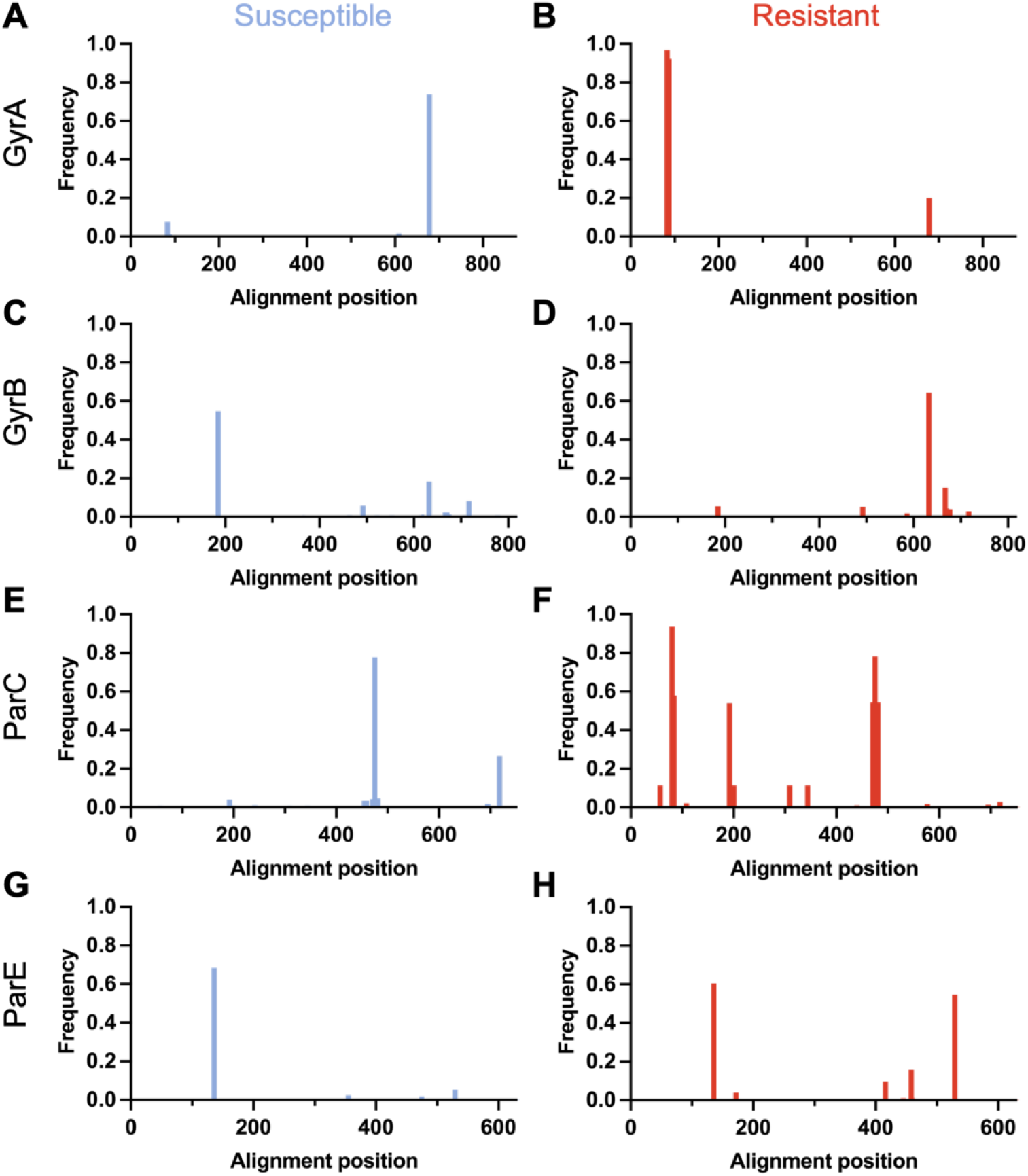
Ciprofloxacin resistance is linked to genetic variation in natural *E. coli* isolates. Genetic variation of ciprofloxacin resistance-related proteins was analysed in a natural population of 1509 *E. coli* isolates. Amino acid changes at each protein alignment position for GyrA (A+B), GyrB (C+D), ParC (E+F), and ParE (G+H), are given in blue or red, to indicate the ciprofloxacin susceptible (n = 1229), and ciprofloxacin resistant (n = 280) subpopulations, respectively.

**Figure 5.**
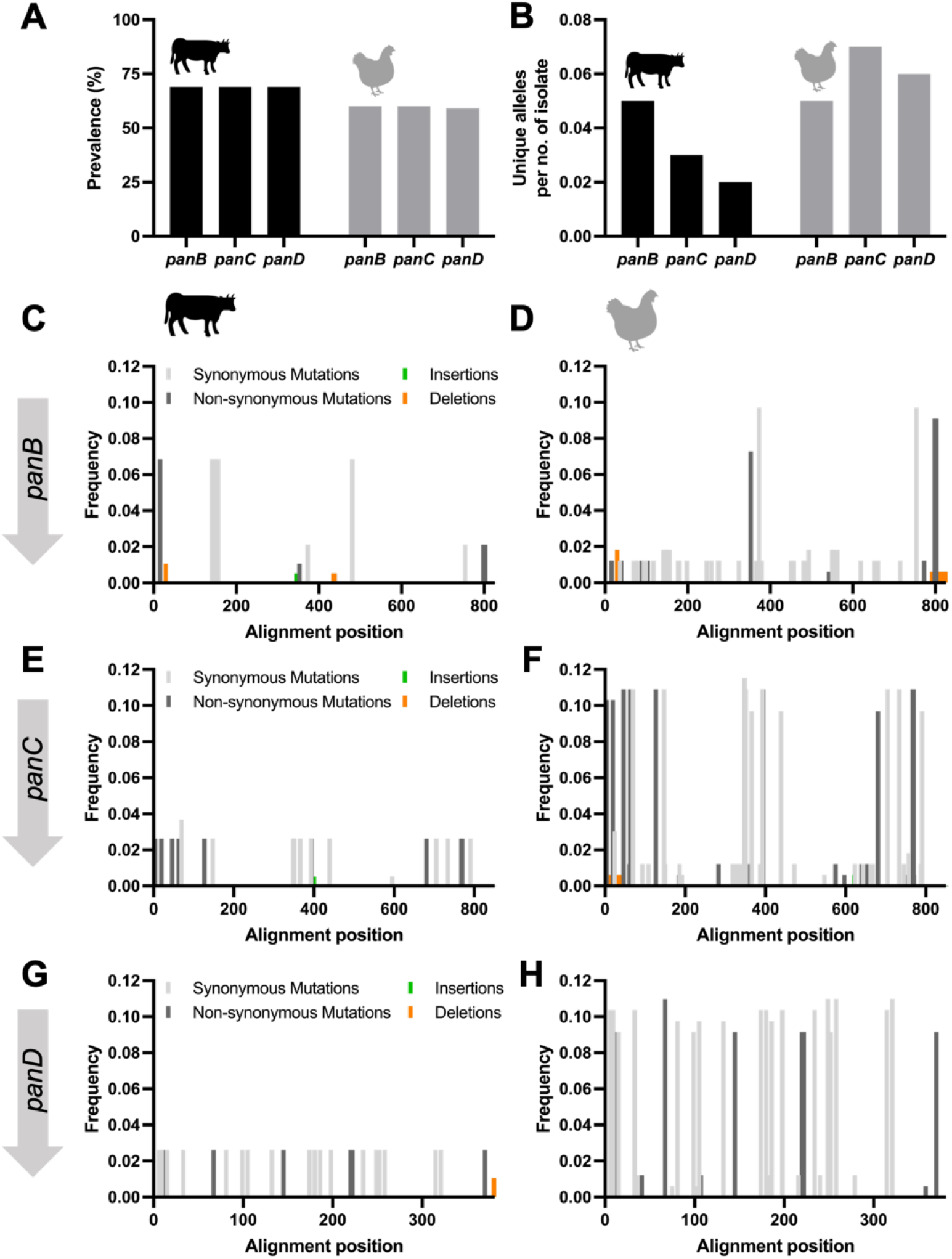
Conservation and diversity of vitamin B5 biosynthesis genes in *Campylobacter jejuni* from cattle and chickens. A: The *panBCD* locus is present in a higher proportion of cattle isolates compared to chicken isolates. B: Chicken-associated isolates harbour a greater number of unique *panBCD* alleles relative to cattle isolates. C-G: Patterns of genetic variation across the *panBCD* locus show reduced diversity in cattle isolates compared with chicken isolates.

**Figure 6.**
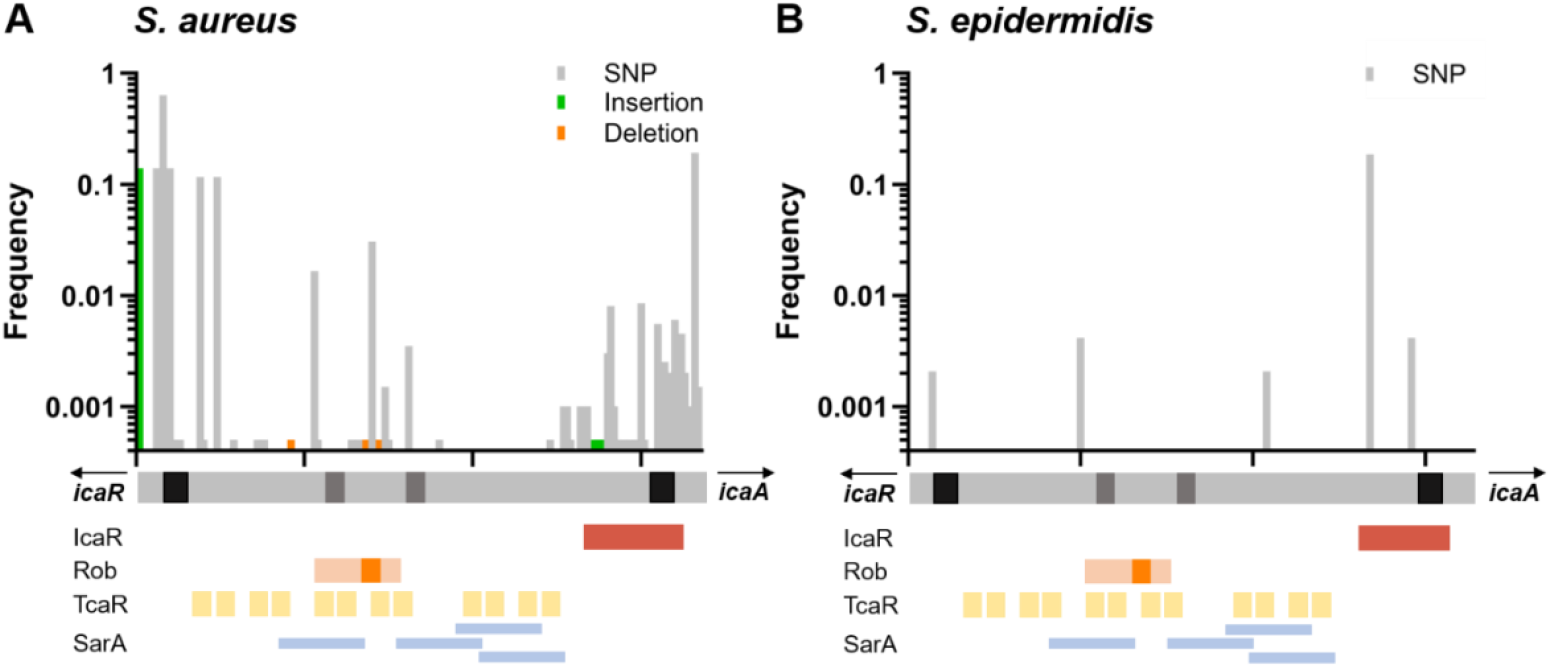
Genetic variation at the non-coding regulatory regions of the biofilm-associated *ica* operon in *S. aureus* and *S. epidermidis*. Single nucleotide polymorphisms (grey), insertions (green) and deletions (orange) are indicated along the nucleotide alignment (grey bar below graph) of the non-coding region between *icaR* and *icaA* in *S. aureus* (A) and *S. epidermidis* (B). Graphical representation of regulatory elements to scale: black: Shine-Dalgarno sequence for *icaR* and *icaA*, respectively; dark grey: likely −35 and −10 regions for transcription initiation of *icaADBC* operon; red: IcaR binding site; orange: Rob binding site with TATTT motif; yellow: TcaR binding sites; blue: SarA binding sites.

### Case study 3: Genetic variation in biofilm-regulating intergenic region in staphylococci

The genome assemblies of *S. aureus* (n=1984), *S. epidermidis* (n=1000), *S. hominis* (n=13), *S. haemolyticus* (n=44), *S. pseudintermedius* (n=178) and *S. chromogenes* (n=49) were accessed on the PubMLST database [20] (Supplementary file 7). Nucleotide sequences of the *icaR*_*icaA* intergenic region of *S. aureus* and *S. epidermidis* were downloaded from GenBank and used to query against isolate assemblies using using blastn (BLAST 2.12.0+ with settings: word size: 20; reward: 2; penalty: −3; gapopen: 5; gapextend: 2). The identified matching sequences were extracted and aligned using MAFFT v7.505 with default parameters to create the nucleotide alignment input file. GeneScanner was run in ‘nucleotide’ mode, selecting strain USA300_FPR3757 (PubMLST *Staphylococcus aureus* ID: 37463) and N13018T (PubMLST *Staphylococcus epidermidis* ID: 41156) as reference for *S. aureus* and *S. epidermidis*, respectively. As the NucleotideAnalysis sheet will still assume a coding sequence, the descriptions *synonymous, non-synonymous* and *stop codons*, should be ignored and mutation frequencies added together. Mutation frequencies were normalised to the isolate number and visualised using GraphPad Prism (version 10.5.0). Information on the regulator binding sites of *icaR*_*icaA* intergenic region was taken from previous studies [50,52–55].

## Supporting information

SupplementaryFile1_syn_nonsyn_alignment

SupplementaryFile2_syn_nonsyn_output

SupplementaryFile3_indel_alignment

SupplementaryFile4_indel_output

SupplementaryFile5_simulation_alignment

SupplementaryFile6_simulation_output

SupplementaryFile7_isolates

## Data availability

GeneScanner code and detailed requirements for each release version are publicly available (https://github.com/Sheppard-Lab/GeneScanner, DOI: 10.5281/zenodo.17495646. GeneScanner requires Python v3.6+. GeneScanner is also available as plugin in the PubMLST online database (https://bigsdb.readthedocs.io/en/latest/data_analysis/genescanner.html). All genome assemblies used are available through the PubMLST database. Isolate IDs, query sequences, and alignment and output files for designed and synthetic datasets can be found in the supplementary files.

## Author contributions

Conceptualization: SKS, CMK

Design & Methodology: SK, PSR, CMK, SKS, BAS

Implementation: SK, PSR, BAS, KJ

Analysis: CMK, SK, PSR, SKS

Data Curation: CMK, SK, PSR, BAS, KJ

Writing - Original Draft: CMK, SK, PSR, SKS

Writing - Review & Editing: all authors

Visualization: CMK, SK

Supervision: SKS, CMK

Project administration: SKS, CMK

Funding acquisition: SKS; CMK

